# Experience Reorganizes Coordinated Population Dynamics Across Hippocampal Circuits

**DOI:** 10.64898/2026.01.15.698118

**Authors:** Shahrukh Khanzada, Xin Hu, Diana Klütsch, Gerd Kempermann, Fabio Boi, Hayder Amin

## Abstract

Hippocampal function relies on structured patterns of population activity that reflect the intrinsic organization of the circuit. Although experience is known to influence synaptic properties and local activity features, whether prolonged experience reshapes the large-scale organization of population dynamics across hippocampal subfields in the intact brain has remained unclear. Progress on this question has been limited by the lack of measurements that capture circuit-wide activity simultaneously while preserving spatial circuit structure. Here, we combined prolonged environmental experience with large-scale, simultaneous multi-shank recordings spanning CA1, CA3, and dentate gyrus in the mouse hippocampus to examine experience-dependent changes in circuit dynamics. We find that long-term experience reorganizes the basal operating state of the hippocampus, altering the statistical structure of field potentials, population spiking, and sharp-wave ripple activity in a coordinated manner across subfields. These changes are not confined to specific regions but instead reflect a circuit-wide reconfiguration of functional interactions that extends over larger spatial scales. At the population level, experience shifts hippocampal dynamics into a more coordinated dynamical regime, characterized by shared low-dimensional structure across hippocampal subfields. Together, these findings demonstrate that long-term experience reorganizes coordinated population dynamics across hippocampal circuits, establishing an intrinsic activity organization that reflects the experiential history of a memory network.

## Main

Experience continuously shapes brain circuits, yet its influence is expressed through the physical substrate by which neural activity is generated, propagated, and integrated within circuits^1,2^. Neural computation does not arise from individual neurons or synapses in isolation but emerges from the collective dynamics of interacting populations, shaped by circuit connectivity, biophysical constraints, and ongoing activity patterns^3–5^. From this perspective, experience-dependent plasticity must ultimately be expressed as changes in circuit-level dynamics: the recruitment of neuronal populations, the spatial organization of interactions, and the temporal structure through which activity propagates and stabilizes^6–8^.

The hippocampus provides a canonical system for studying experience-dependent circuit organization. Its trisynaptic architecture, recurrent and feedforward motifs, and well-characterized oscillatory dynamics support functions ranging from memory encoding to consolidation and planning^9–11^. Experience modifies hippocampal circuits at multiple levels, including synaptic strength, intrinsic excitability, and structural connectivity^2,12,13^. Environmental enrichment, in particular, has been shown to enhance synaptic plasticity, alter oscillatory structure, modulate sharp-wave ripple activity and functional connectivity, and improve learning and memory^14–18^. However, these findings have largely been derived from sparse recordings or localized *ex vivo* measurements, leaving unresolved how experience reorganizes interactions across intact hippocampal circuits.

A growing body of theoretical and experimental work emphasizes that circuit computation depends on population-level properties, such as coordination geometry, dimensionality, and shared activity modes^19–23^. These properties determine how information is represented, transmitted, and transformed, and they cannot be inferred from isolated neurons or single regions. In the hippocampus, transient events such as sharp-wave ripples provide a clear example: their functional role depends not only on local generation but also on coordinated recruitment and propagation across CA3, CA1, and the dentate gyrus (DG)^24–26^. Yet capturing such coordination *in vivo* requires simultaneous, dense sampling across multiple subfields and layers—capabilities that have remained technically limited.

This limitation is fundamentally technological. Conventional electrophysiological approaches sample only a small fraction of the hippocampal network, while optical methods lack the temporal resolution needed to resolve fast population events and their coordination. As a result, it remains unclear how experience modulates basal operating states, interaction geometry, event structure, and population dimensionality within the intact hippocampus. These features are increasingly recognized as central determinants of neural computation, but they have been challenging to access simultaneously *in vivo*^27,28^.

We previously addressed aspects of this problem *ex vivo* using large-scale CMOS-based recordings, demonstrating that environmental enrichment induces persistent changes in hippocampal connectivity and network dynamics at the circuit level^15,16,29–31^. While *ex vivo* preparations enable dense spatial sampling and precise control, they necessarily disrupt long-range coordination, neuromodulatory influences, and ongoing brain state. Whether and how experience-dependent circuit reorganization manifests in the intact brain—where computation unfolds across distributed, interacting populations—has remained an open question.

Here, we resolve this gap by combining prolonged environmental enrichment with high-density, large-scale *in vivo* electrophysiological recordings spanning the hippocampal formation. Using an active CMOS SiNAPS probe that simultaneously records from 1024 electrode-pixels^32^ across multiple hippocampal subfields, we examine experience-dependent changes in the basal network state, functional interaction geometry, ripple organization, and population-level dynamics within a unified analytical framework. By treating experience as a factor that reorganizes circuit-level dynamics rather than as a modulation of isolated signals, this study provides a principled account of how experiential history reconfigures hippocampal computation *in vivo*.

Beyond hippocampal physiology, these results have broader implications for the design of neural interfaces and adaptive neuromodulation strategies. Effective interaction with neural systems—whether for decoding, stimulation, or the restoration of function—requires understanding how experience structures population dynamics and limits the space of accessible neural states^33,34^. By revealing how experience reorganizes circuit-level dynamics in the intact brain, this work defines empirically grounded constraints relevant to computational models, closed-loop interfaces, and future circuit-based intervention strategies.

## Results

### High-density Electroanatomical Recordings Enable *in vivo* Interrogation of Experience-dependent Hippocampal Computation

Understanding how experience reshapes hippocampal computation requires simultaneous access to neuronal population activity, mesoscopic field dynamics, and anatomical context across distributed circuit elements in the intact brain. To meet this challenge *in vivo*, we combined prolonged environmental enrichment with large-scale, high-density electrophysiological recordings spanning the transverse axis of the hippocampal formation (**Fig. 1a, c**).

**Figure 1.**
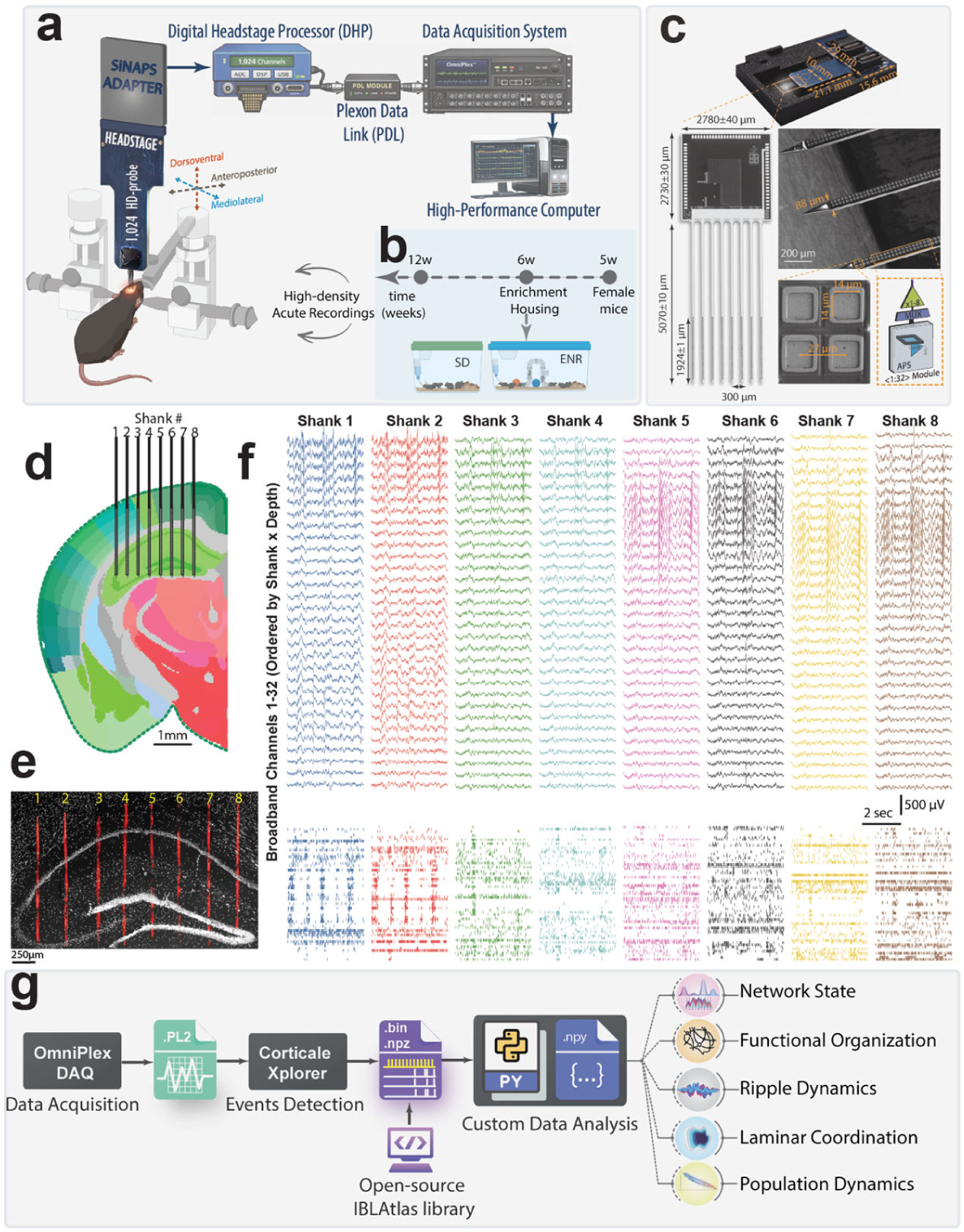
Experimental framework and high-density electroanatomical recording strategy. a, Stereotaxic implantation strategy showing probe trajectory and orientation relative to the hippocampal formation, designed to span multiple hippocampal subfields along the transverse axis. **b,** Experimental design. Mice were housed under standard (SD) or enriched (ENR) conditions for ≥6 weeks before *in vivo* recordings to induce sustained experience-dependent circuit adaptations. **c,** Eight-shank active CMOS SiNAPS probe, which integrates 1024 simultaneously active electrode-pixels distributed across 8 parallel shanks (128 channels per shank) with 300 µm inter-shank spacing, enabling continuous two-dimensional sampling across hippocampal circuitry. Each shank comprises modular front-end units with platinum electrode-pixels (∼25–30 µm pitch), on-pixel amplification, filtering, and time-division multiplexing, allowing full-bandwidth acquisition from all channels without channel selection or temporal subsampling. **d,** Histological slice with overlaid shank positions used to identify the precise anatomical location of each shank across CA1, CA3, and DG. This panel serves as the reference for electroanatomical assignment of recording channels. **e,** Post hoc histological reconstruction with (DAPI; grey) and (CM-DiI; red) confirming consistent probe placement across hippocampal subfields and laminar domains, with individual shanks traversing principal cell layers and dendritic laminae. **f,** Representative broadband recordings acquired simultaneously across shanks and depths, showing concurrent capture of local field potentials, sharp-wave ripple events, and high-frequency population spiking within the same trials. **g,** Hardware–analysis integration workflow. Dense electroanatomical recordings are combined with atlas-based registration and custom computational pipelines to enable joint analysis of laminar dynamics, inter-regional interactions, event structure, and population-level activity within a unified circuit reference frame.

Mice were housed either under standard laboratory conditions (SD) or in an enriched environment (ENR) for a minimum of six weeks, a duration sufficient to induce sustained experience-dependent plasticity rather than transient behavioral modulation (**Fig. 1b**). While we previously demonstrated *ex vivo* that environmental enrichment induces lasting changes in hippocampal connectivity and network dynamics across scales^15,16^, establishing how these alterations are expressed at the level of distributed circuit computation in the intact brain has remained technically challenging. In vivo, such questions require simultaneous access to population spiking, mesoscopic field dynamics, and their anatomical organization across hippocampal subfields—capabilities that have been limited by sparse spatial sampling and restricted channel counts.

To overcome these limitations, we implemented high-density electroanatomical recordings using a multi-shank active CMOS SiNAPS probe embedded within a fully integrated, custom-built *in vivo* recording framework. The probe integrates 1024 simultaneously active electrode-pixels distributed across eight parallel shanks with 300 µm inter-shank spacing, enabling continuous two-dimensional sampling over more than 4 mm² of tissue (**Fig. 1c**). This configuration permits concurrent access to laminar, inter-regional, and population-scale activity within a single recording, preserving circuit geometry while maintaining full bandwidth acquisition across all channels. Each shank comprises modular front-end units with closely spaced platinum electrode-pixels and on-pixel amplification and multiplexing, allowing concurrent acquisition of local field potentials, sharp-wave ripple activity, and high-frequency spiking signals from all channels without temporal subsampling or channel selection^32^. This architecture differs fundamentally from conventional linear or partially multiplexed probes by preserving full spatial resolution across the entire array during single recordings.

Probes were implanted stereotaxically to span CA1, CA3, and DG along the hippocampal transverse axis (**Fig. 1d**). Post hoc histological reconstruction confirmed consistent probe placement across animals, with individual shanks traversing defined hippocampal layers across CA1, CA3, and DG, including principal cell layers and their associated dendritic laminae (**Fig. 1e**). This configuration enabled simultaneous recording from multiple hippocampal subfields and layers, providing an electroanatomical view of circuit dynamics that is inaccessible with single-shank or region-restricted recordings. Broadband recordings obtained across the full array revealed the simultaneous presence of mesoscopic field dynamics, sharp-wave ripple activity, and distributed population spiking within the same trials and anatomical context (**Fig. 1f**). Crucially, this simultaneous access enables direct comparison of how experience-dependent modifications are expressed across signals that reflect synaptic integration, neuronal output, and coordinated network events, rather than inferring such relationships across separate experiments or preparations. The resulting dataset permits analyses that explicitly bridge organizational scales, enabling the examination of field dynamics, population spiking, laminar structure, and inter-regional interactions within a single anatomical and temporal framework. Rather than treating these signals as separate modalities, the analysis pipeline integrates them at the level of circuit computation, allowing relationships between local activity patterns and distributed network dynamics to be resolved directly (**Fig. 1g**). This hardware–analysis co-design establishes an experimental regime in which experience-dependent changes can be studied as transformations of circuit-level computation in the intact brain. By maintaining continuous spatial coverage and preserving population structure across recording and analysis stages, the framework enables interrogation of how experience reshapes coordinated activity patterns across hippocampal subfields—an essential prerequisite for future strategies aimed at selectively interfacing with and manipulating circuit-specific dynamics *in vivo*.

### Basal Hippocampal Network State is Restructured by Experience

To determine whether prolonged environmental enrichment alters the intrinsic operating regime of the hippocampus in the intact brain, we quantified the basal activity state of hippocampal networks during spontaneous activity. Basal state here refers to the joint organization of mesoscopic field dynamics, population spiking output, and structured network events, which together define how the circuit operates in the absence of explicit task demands.

We first examined local field potential (LFP) activity, which reflects the collective synaptic and transmembrane currents integrating across local neuronal populations and provides a mesoscopic readout of network engagement^27,35^. Depth-and shank-resolved electroanatomical maps revealed that enriched animals exhibited a markedly altered spatial organization of LFP activity across CA1, CA3, and DG compared to standard-housed controls (**Fig. 2a**). Rather than a uniform scaling, enrichment engaged a broader fraction of the hippocampal volume, indicating a global shift in baseline network engagement.

**Figure 2.**
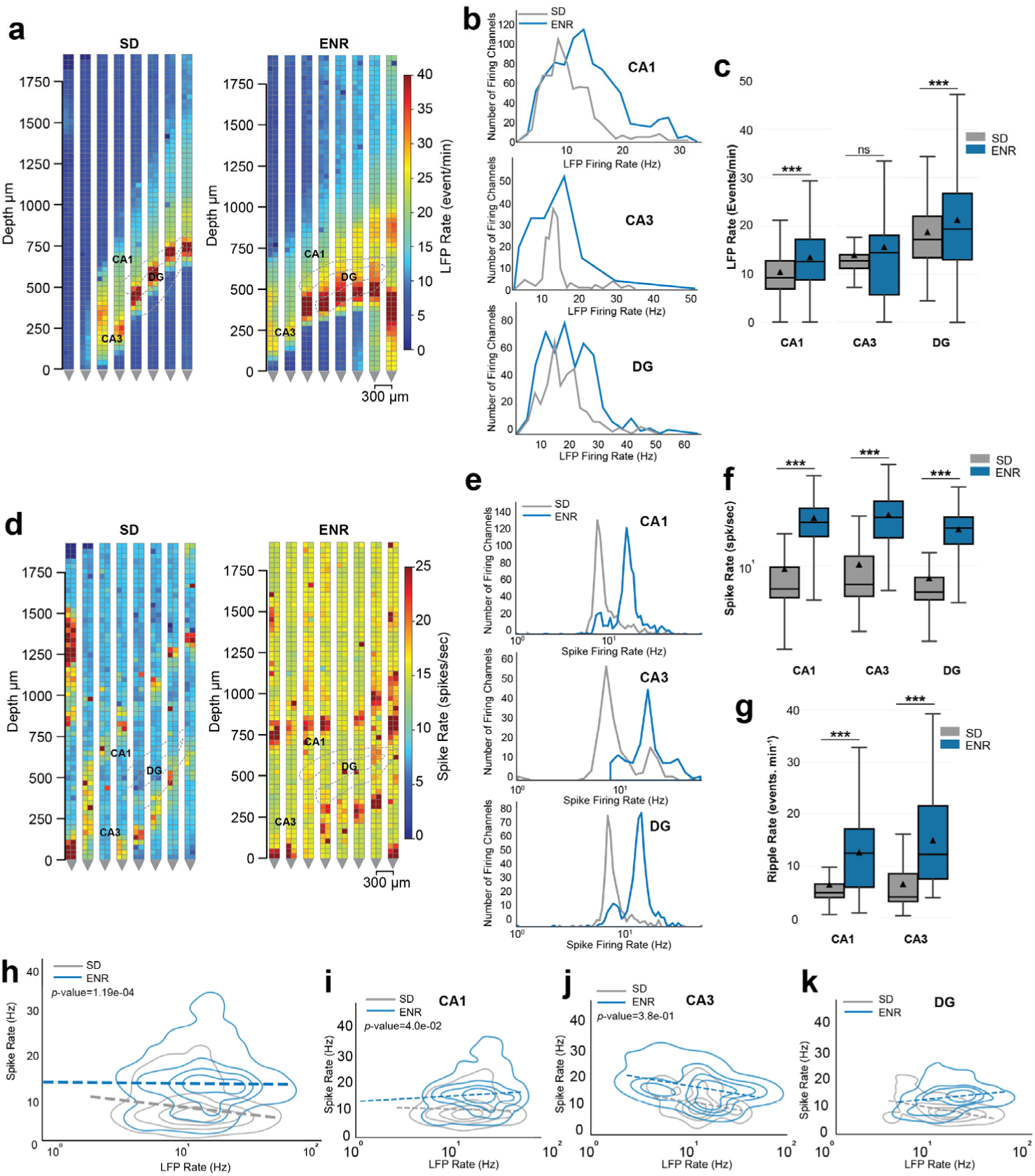
Baseline hippocampal field, spiking, and SWR activity across subfields. a, Depth-and shank-resolved electroanatomical maps of broadband LFP activity across CA1, CA3, and DG in SD and ENR animals, revealing a broader spatial engagement of the hippocampal network in ENR. **b,** Distributions of instantaneous LFP rate plotted on logarithmic scales for CA1, CA3, and DG, showing a rightward shift toward larger-amplitude events in ENR. *(CA1, p*<10^-24^, CA3 *p*<0.05, DG *p*<10^-4^ *Kolmogorov–Smirnov test*) **c,** Region-averaged LFP activity rates in SD and ENR animals. (\*\*\**p* < 0.001 *ANOVA*). **d,** Channel-level maps of multiunit firing activity across shanks and depths for SD and ENR recordings. **e,** Distributions of channel-level firing rates for CA1, CA3, and dentate gyrus, demonstrating a shift toward higher firing activity in ENR. *(CA1, p*<10^-78^, CA3 *p*<10^-21^, DG *p*<10^-67^ *Kolmogorov–Smirnov test*). **f,** Mean population firing rates per region in SD and ENR animals. (\*\*\**p* < 0.001 *ANOVA*). **g,** SWR occurrence rates in CA3 and CA1 for SD and ENR animals. (\*\*\**p* < 0.001 *ANOVA*). **h–k,** Relationship between broadband LFP activity and multiunit firing rate across channels for all hippocampal regions, illustrating strengthened spike–field coupling in ENR.

At the level of activity distributions, LFP amplitudes displayed a pronounced heavy-tailed structure across all regions (**Fig. 2b**). In enriched animals, channel-wise activity distributions were shifted toward higher firing rates on logarithmic scales, with a larger fraction of channels contributing to high-activity regimes. This redistribution reflects a reweighting of population activity toward rare but strongly expressed network states, consistent with heavy-tailed statistics observed in large neuronal populations, where collective dynamics are dominated by infrequent, high-impact events rather than Gaussian fluctuations^36,37^. Consistent with this interpretation, region-averaged LFP activity rates were significantly elevated in enriched animals across CA1, CA3, and DG (**Fig. 2c**).

We next asked how this altered mesoscopic regime is reflected in population spiking output, which represents the final expression of network activity. Channel-level multiunit activity (MUA) firing maps demonstrated higher firing activity across shanks and depths in enriched animals (**Fig. 2d**). As with LFPs, MUA firing-rate distributions were shifted toward higher values in enrichment across all hippocampal regions (**Fig. 2e**), resulting in elevated mean population firing rates in CA1, CA3, and DG (**Fig. 2f**). Importantly, these effects were observed at the level of population activity recorded by dense electrode arrays, without relying on single-unit isolation, consistent with prior observations that hippocampal firing statistics are intrinsically skewed and scale with network state^38^.

Further, we assessed the incidence of sharp-wave ripple (SWR) events, the most prominent intrinsic coordination pattern of the hippocampus, and a canonical signature of internally generated hippocampal communication^25,39^. Ripple occurrence was significantly elevated in enriched animals in both CA3 and CA1 (**Fig. 2g**), indicating that enrichment increases the spontaneous expression of highly coordinated network events central to hippocampal information processing.

To directly link mesoscopic and microscopic dynamics, we quantified the relationship between broadband LFP activity and population spiking. Enriched animals exhibited a more substantial alignment between LFP power and MUA spiking output across channels and regions (**Fig. 2h–k**), indicating that enrichment tightens the coupling between synaptic-scale network activity and neuronal output across the hippocampal circuit. Such strengthened spike–field coupling reflects tighter coordination between synaptic population activity and neuronal output across hippocampal subfields^35,40^.

Together, these findings demonstrate that environmental enrichment places the hippocampus into a distinct basal operating regime *in vivo*, characterized by enhanced network engagement, heavier-tailed activity distributions, strengthened spike–field coupling, and increased expression of coordinated ripple events.

### Experience-dependent Scaling of Spatial Organization and Topology of Hippocampal Functional Interactions

Having established that environmental enrichment alters the basal activity regime of the hippocampus *in vivo* (**Fig. 2**), we next asked whether this shift is accompanied by a reorganization of functional interactions coordinated across space and across hippocampal subfields. Functional interactions, quantified as correlated activity across space and regions, regulate information flow in the hippocampus and determine whether activity remains locally confined or is distributed across the circuit, thereby serving as a critical bridge between the basal state and structured network events. It provides a direct measure of network integration beyond changes in activity magnitude alone.

We first examined LFP-based functional connectivity across hippocampal subfields. Region-by-region connectivity matrices revealed a distinct increase in correlated LFP activity in enriched animals compared to standard-housed controls, both within individual regions and across CA3–CA1–DG interactions (**Fig. 3a**). This enhancement was not restricted to a single pathway but reflected a global strengthening of mesoscopic coupling across the hippocampal network. Quantification at the animal level confirmed that LFP correlations between all major region pairs (CA3–CA1, CA1–DG, CA3–DG) were systematically higher in enriched animals (**Fig. 3b**), indicating that enrichment promotes coordinated activity across anatomically distinct hippocampal subcircuits. Given the distinct anatomical and computational roles of hippocampal subfields, such cross-regional strengthening indicates a circuit-level reorganization rather than a uniform increase in local synchrony^41,42^.

**Figure 3.**
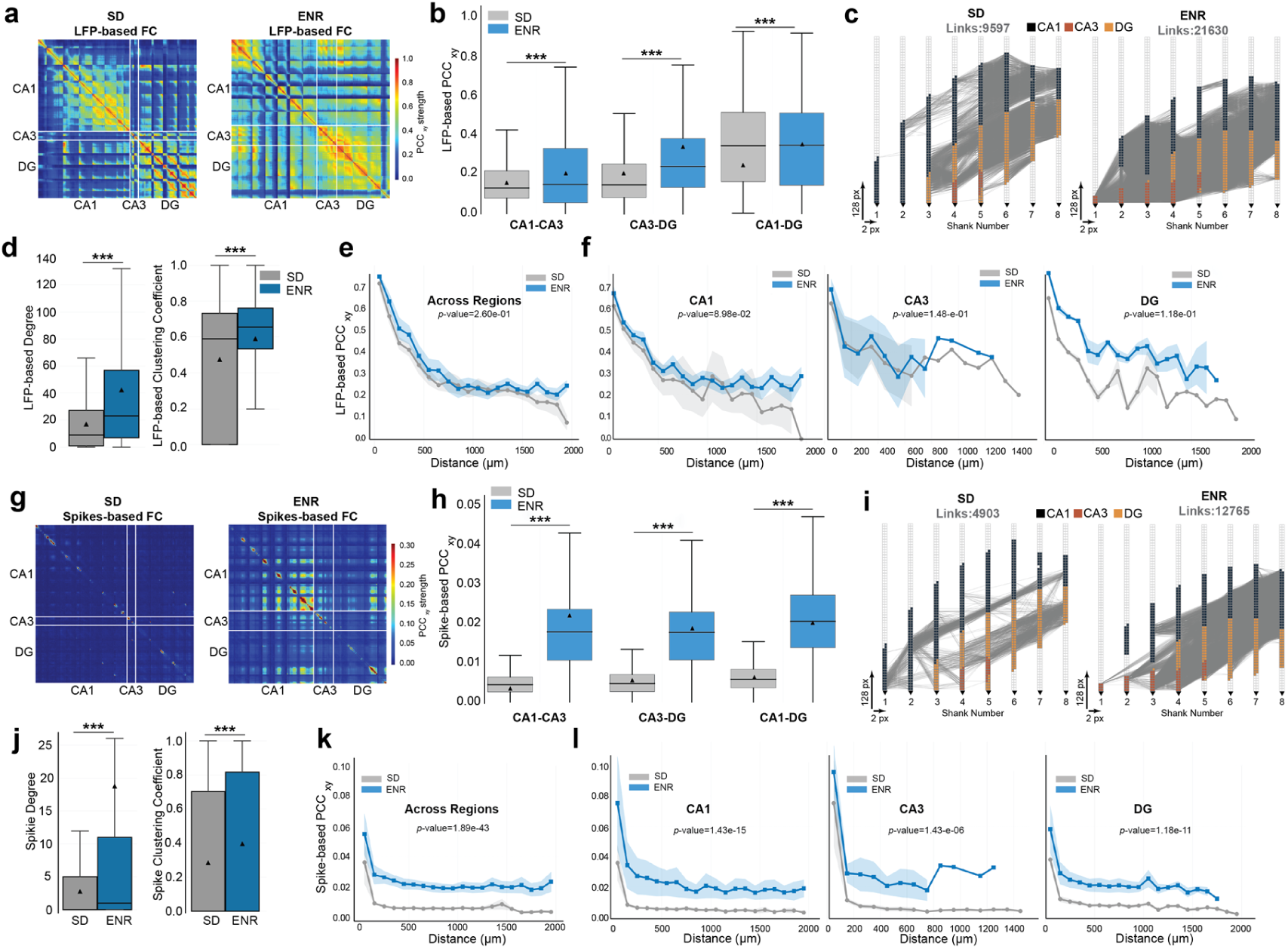
Spatial organization and distance dependence of hippocampal functional connectivity. a, Region-by-region LFP functional connectivity matrices for SD ENR animals, showing increased within-and cross-region correlations in ENR across CA1, CA3, and DG. **b,** Animal-level quantification of LFP connectivity for key hippocampal region pairs (CA3–CA1, CA1–DG, CA3–DG), revealing stronger coupling in ENR. **c,** LFP-based functional connectivity graphs mapped onto the 1024-channel shank geometry for SD and ENR recordings, illustrating denser and more spatially extensive interaction patterns in ENR. **d,** Graph-theoretic metrics (mean degree and clustering coefficient) for LFP-based networks, demonstrating increased network integration in ENR. **e,** LFP correlation as a function of inter-electrode distance pooled across regions and animals, showing enhanced long-range coordination in ENR. **f,** Region-specific LFP correlation–distance relationships for CA1, CA3, and DG. **g–h,** Spike-based region-by-region functional connectivity matrices and animal-level statistics, revealing enhanced spiking correlations in ENR. **i–j,** Spike-based functional connectivity graphs and corresponding graph metrics, demonstrating increased network density and clustering in ENR. **k–l,** Spike correlation as a function of distance across and within hippocampal regions, indicating extended spatial coordination in ENR.

To visualize how these interaction changes are embedded in physical space, we reconstructed LFP-based functional connectivity graphs using the full 1024-channel geometry. In enriched animals, connectivity maps revealed a denser and more spatially extensive interaction structure, with numerous long-range links spanning shanks and hippocampal regions, in contrast to the sparser and more locally confined networks observed in standard-housed animals (**Fig. 3c**). This spatial embedding reveals that enrichment reorganizes the hippocampus into a more globally connected interaction network, rather than merely strengthening existing local couplings.

Consistent with these qualitative observations, graph-theoretic metrics demonstrated significantly higher node degree and clustering coefficient in enriched networks (**Fig. 3d**), indicating that enrichment increases both the number of functional connections and their local organization into interconnected clusters. Such changes in network topology reflect altered interaction structure and communication efficiency within functional brain networks^16,43,44^.

Beyond network topology, we next asked whether experience-dependent circuit reorganization alters the spatial scale over which hippocampal activity is coordinated. Analysis of LFP correlation as a function of inter-electrode distance revealed that enriched animals exhibited higher correlations across both short and long spatial scales (**Fig. 3e**). Importantly, this enhancement persisted at larger distances, demonstrating that enrichment extends the range over which coherent mesoscopic dynamics are expressed. Region-specific analyses confirmed that this effect was present within individual hippocampal subfields, including CA1, CA3, and DG (**Fig. 3f**), indicating that enrichment increases long-range coordination both locally and globally. Distance-dependent correlation structure directly reflects the effective range of functional interactions and suggests that enrichment expands the spatial extent over which coherent hippocampal dynamics are expressed^45,46^.

To determine whether this reorganization of interaction structure is also reflected in neuronal output, we implemented similar analyses using channel-level multiunit spike correlations. As observed for LFPs, enriched animals exhibited stronger spike-based functional connectivity across hippocampal regions (**Fig. 3g–h**), denser spike correlation graphs with extensive cross-regional links (**Fig. 3i**), and significantly elevated graph metrics (**Fig. 3j**). Spike correlation also decayed more slowly with distance in enriched animals, both across regions and within individual subfields (**Fig. 3k–l**), demonstrating that enrichment enhances the spatial extent of coordinated population output.

Together, these results demonstrate that ENR reorganizes the hippocampal network into a more integrated and spatially extended interaction regime, strengthening functional coupling within and across subfields and aligning field and spiking dynamics across the circuit.

### Experience-dependent Modulation of Ripple Structure and Circuit Recruitment

Having shown that environmental enrichment shifts the basal activity regime of the hippocampus (**Fig. 2**) and expands the spatial scale and topology of functional interactions across subfields (**Fig. 3**), we next asked how this reconfigured network state manifests during SWR events—the most prominent intrinsically generated coordination pattern of the hippocampus^25,47^. Ripples provide a canonical window into hippocampal circuit dynamics, as they reflect the rapid, synchronous engagement of CA3–CA1 networks and are tightly linked to hippocampal communication and memory-related processing^48,49^. We first examined individual ripple events across the full multi-shank geometry to assess whether enrichment alters ripple structure at the single-event level. Representative ripple-band traces revealed salient qualitative differences between SD and ENR animals (**Fig. 4a**). In enriched animals, ripple events exhibited higher amplitude and broader engagement across shanks compared to the more spatially confined, lower amplitude ripples observed in SD. Region-resolved ripple waveforms from CA3 and CA1 further demonstrated that enrichment alters ripple signatures themselves, increasing event amplitude and producing more sharply defined ripple waveforms compared to standard housing (**Fig. 4b**). These examples indicate that enrichment does not merely change how often ripples occur, but alters their intrinsic structure.

**Figure 4.**
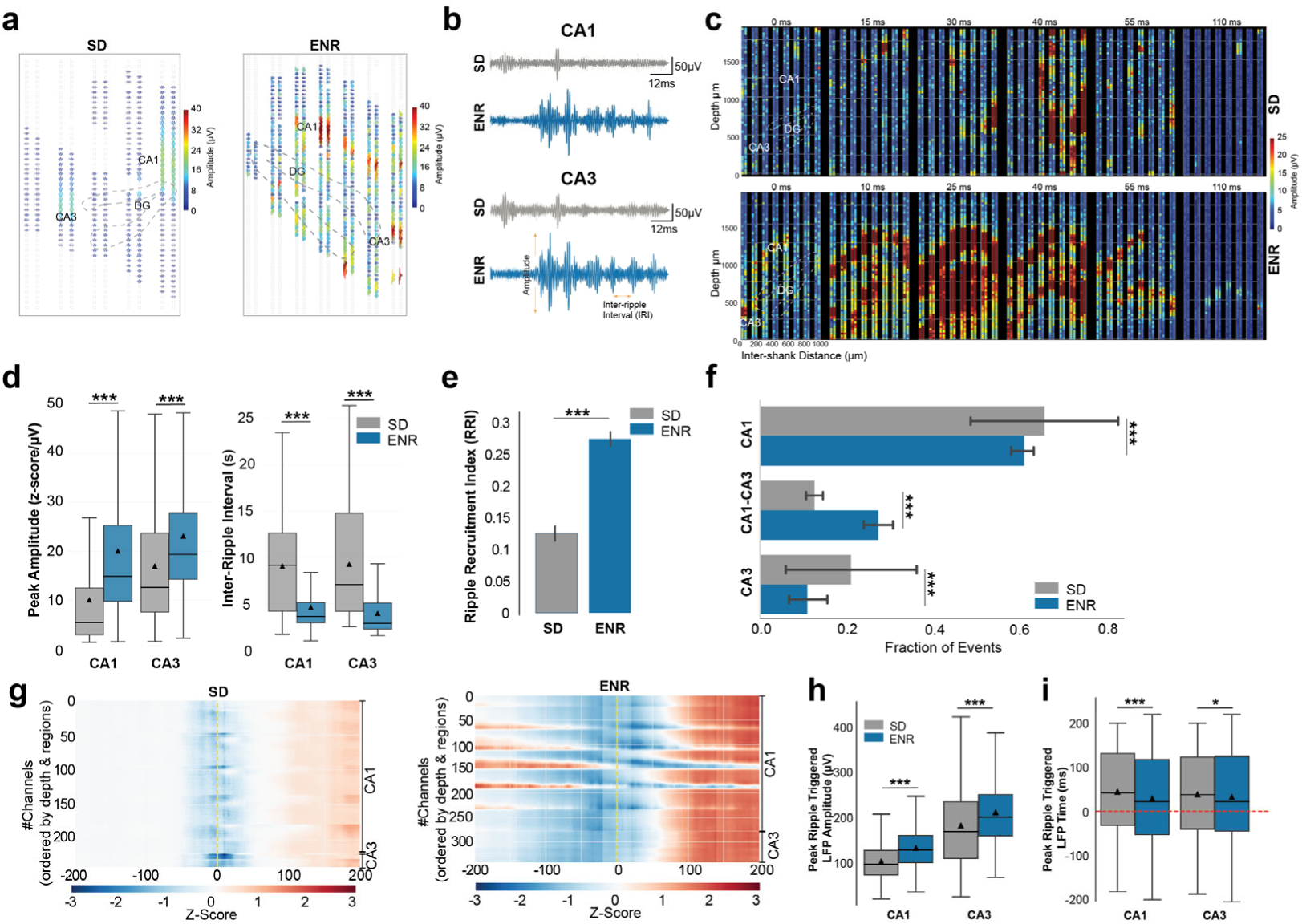
Spatiotemporal structure and circuit recruitment of hippocampal ripple events a, Example ripple-band traces recorded across multiple shanks and depths during individual ripple events, illustrating differences in spatial engagement between SD and ENR. **b.** Region-resolved ripple waveforms from CA3 and CA1, showing condition-dependent changes in ripple amplitude and temporal waveform structure. **c,** Spatiotemporal maps of ripple-band amplitude across channels (shank × depth) for representative ripple events, revealing differences in the spatial footprint and temporal evolution of ripple activity. **d,** Quantification of ripple peak amplitude and inter-ripple interval in CA3 and CA1, demonstrating differences in ripple strength and event density across conditions. **e,** RRI, measuring the fraction of ripple events that co-engage CA3 and CA1, indicating changes in inter-regional ripple propagation. **f,** Distribution of channel-level recruitment during ripple events, showing the proportion of ripples confined to single regions versus those engaging both CA3 and CA1. **g,** Ripple-triggered averages of broadband LFP activity across depth and region, illustrating laminar circuit dynamics associated with ripple initiation and propagation. **h,** Peak amplitude of ripple-triggered LFP responses in CA3 and CA1. **i,** Latency of ripple-triggered LFP peak relative to ripple onset, reflecting the temporal coordination of laminar circuit engagement.

To visualize how ripple activity unfolds across space and time^50,51^, we constructed two-dimensional spatiotemporal maps of ripple-band amplitude across channels and shanks (**Fig. 4c**). In SD animals, ripple activity emerged as a narrow, focal hotspot localized near its site of origin. In contrast, ENR animals exhibited ripple events with a broader spatial footprint and more pronounced temporal evolution, indicating that ripple activity spreads across larger portions of the hippocampal network as it develops.

We next quantified these observations across all detected ripple events and animals. Ripple peak amplitudes were significantly higher in enriched animals in both CA1 and CA3, while inter-ripple intervals were shorter (**Fig. 4d**), demonstrating that enrichment increases both the strength and temporal density of ripple events. However, amplitude and rate alone do not capture the extent to which ripples engage the hippocampal circuit. To directly address spatial recruitment, we quantified whether individual ripples remained confined to their site of origin or propagated across subfields.

We defined a Ripple Recruitment Index (RRI) that captures the fraction of ripple events engaging both CA3 and CA1 relative to those remaining local^4,52^. Enriched animals exhibited a significantly higher RRI compared to controls (**Fig. 4e**), indicating that enrichment increases the probability that ripples propagate across hippocampal subfields rather than remaining spatially segregated. This shift from local to bi-regional events demonstrates that enrichment promotes large-scale coordination during ripples, consistent with the expanded interaction geometry identified in **Fig. 3**.

To further quantify spatial engagement at the channel level, we measured the fraction of recording sites exceeding a ripple-band threshold during each event (**Fig. 4f**). Enrichment reduced the fraction of ripples confined to CA1-only or CA3-only recruitment, while increasing the fraction of events engaging both regions simultaneously. Importantly, this effect reflects redistribution of ripple propagation rather than suppression of ripple generation, indicating that enrichment converts locally confined ripples into spatially recruited events.

Finally, we examined how enrichment alters the laminar circuit dynamics accompanying ripples^53,54^ by computing ripple-triggered averages of broadband LFP activity across depth and region (**Fig. 4g**). In SD animals, ripple-triggered LFP profiles were characterized by focal, rapidly decaying laminar signatures largely confined to the ripple origin layer. In contrast, ENR animals exhibited pronounced pre-ripple buildup, stronger and vertically extended laminar responses at ripple onset, and sustained post-ripple activity spanning multiple layers and regions. These differences indicate that enrichment enhances both the strength and temporal coordination of laminar circuit engagement during ripple events.

Quantitative features extracted from the ripple-triggered LFP profiles confirmed these observations. ENR animals showed significantly larger ripple-triggered LFP amplitudes (**Fig. 4h**) and shorter peak latencies relative to ripple onset (**Fig. 4i**) in both CA1 and CA3. Because these measures are derived from the same laminar response, their co-variation indicates that enrichment strengthens and accelerates laminar circuit recruitment once a ripple is initiated.

Together, these results demonstrate that environmental enrichment transforms hippocampal ripples from spatially confined events into stronger, faster, and more widely propagating network phenomena, revealing how enrichment-induced changes in the basal state and interaction geometry are expressed during a canonical hippocampal coordination event^24,40^.

### Experience-dependent Convergence of Hippocampal Population Dynamics onto a Coordinated Low-dimensional Regime

The reorganization of basal activity statistics (**Fig. 2**), interaction geometry (**Fig. 3**), and ripple structure (**Fig. 4**) suggests that environmental experience may ultimately restructure hippocampal function at the level of population coding. Population-level state-space analyses provide a principled framework for understanding how large neuronal ensembles jointly evolve, integrating local circuit mechanisms into global dynamical states^22,23^.

We therefore examined region-resolved population dynamics using state-space representations derived from channel-level MUA in CA1, CA3, and DG. Full population trajectories revealed marked differences in the temporal organization of population activity between conditions (**Fig. 5a, left**). In SD animals, trajectories exhibited irregular, multi-directional wandering in state space, with no dominant axis governing temporal evolution. In contrast, ENR animals displayed highly structured trajectories characterized by repeated, directed sweeps along a dominant axis, indicating that population activity evolves through a constrained set of coordinated states. Consistent with this observation, the average population trajectory preserved a pronounced elongated geometry in enriched animals, whereas averaging collapsed trajectories toward the origin in controls (**Fig. 5a, right**), indicating the presence of a stable, shared population mode.

**Figure 5.**
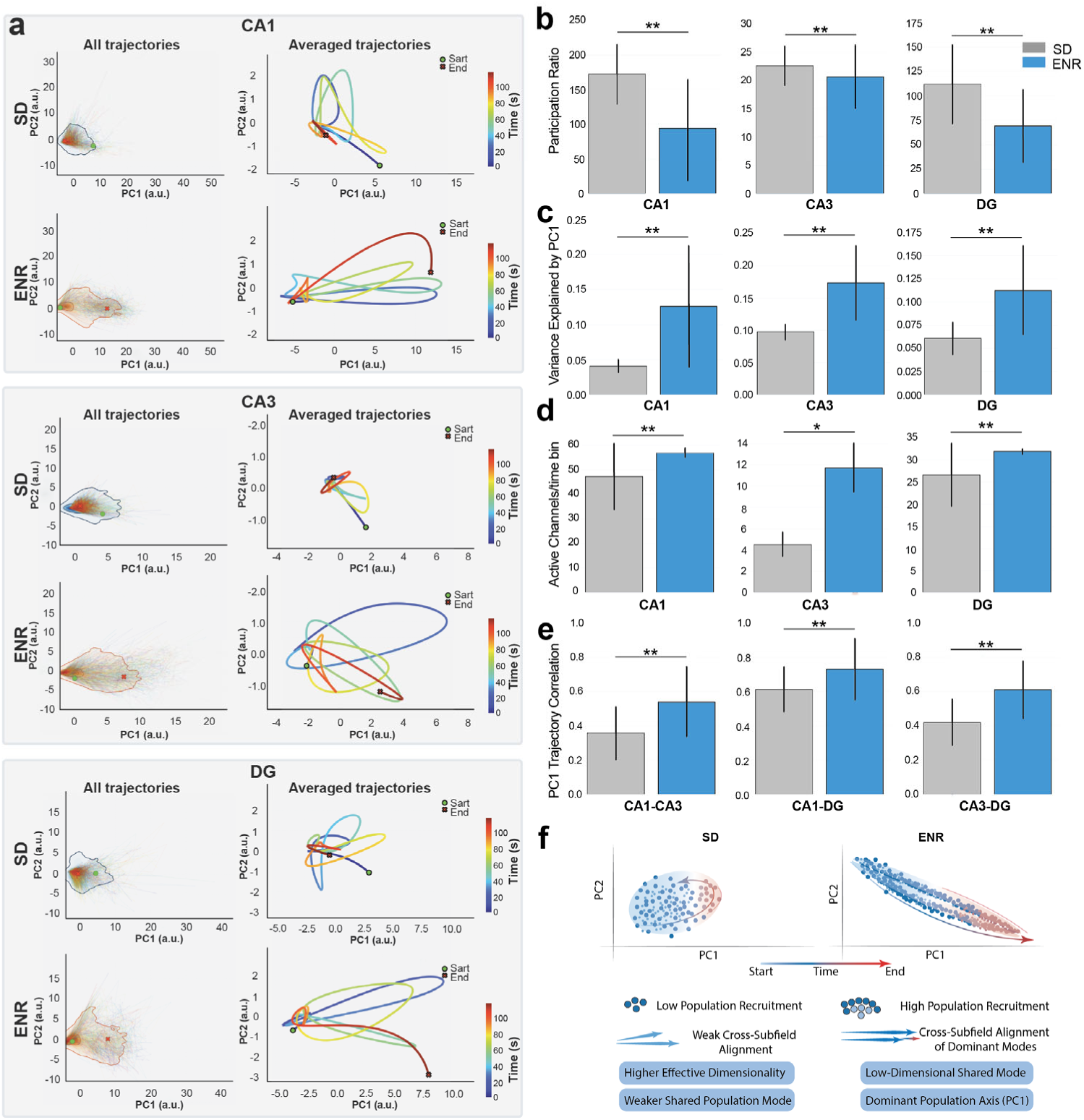
Population trajectories, dimensionality, recruitment, and inter-regional coordination. a, Population state-space trajectories derived from channel-level MUA in CA1, CA3, and DG. *Left* panels show full-time-resolved trajectories projected onto the first two principal components, with color indicating temporal progression. *Right* panels show the corresponding average trajectory over time. ENR animals exhibit recurrent, directed sweeps along a dominant axis, whereas standard-housed animals show more diffuse and irregular trajectories. **b,** Effective dimensionality of population activity quantified by the participation ratio of principal component eigenvalues for CA1, CA3, and dentate gyrus. ENR animals display reduced effective dimensionality across all regions. **c,** Variance explained by the first principal component (PC1 dominance) for each hippocampal subfield, demonstrating strengthened shared population modes in enriched animals. **d,** Population recruitment measured as the number of active channels per time bin within each region. ENR animals recruit a larger fraction of recording sites, indicating coordinated co-activation rather than sparsification. **e,** Inter-regional coordination of population dynamics is quantified as correlations between dominant population trajectories (PC1 time series) across hippocampal subfields, revealing enhanced cross-region alignment in enriched animals. **f,** Conceptual schematic summarizing the population-dynamical regimes identified in Fig. 5a–e. SD animals exhibit diffuse, higher-dimensional population activity without a dominant shared mode, whereas ENR animals exhibit low-dimensional population dynamics dominated by a strong PC1-aligned shared mode, accompanied by broad recruitment and cross-regional alignment.

To quantify these geometric differences, we measured the effective dimensionality of population activity using the participation ratio of principal component eigenvalues. Across CA1, CA3, and DG, enriched animals exhibited significantly lower participation ratios than controls (**Fig. 5b**), demonstrating that population activity occupies a reduced effective dimensionality. This reduction was accompanied by a substantial increase in first-component dominance, with the leading principal component accounting for a larger fraction of total variance in enriched animals across all regions (**Fig. 5c**). Such dominance of shared population modes has been repeatedly observed in hippocampus and other cortical systems and reflects coordinated population activity rather than independent firing across neurons^19,20,38^.

Critically, reduced dimensionality can arise either from diminished activity or from enhanced coordination^55^. To distinguish between these possibilities, we directly quantified population recruitment by measuring the number of active channels per time bin. ENR animals consistently recruited a larger fraction of recording sites across all hippocampal subfields (**Fig. 5d**), indicating that dimensionality reduction reflects compression via co-activation rather than sparsification^28^. Similar reductions in dimensionality accompanying increased recruitment have been described during coordinated brain states and are associated with transitions into integrated population regimes^56–58^. Finally, we asked whether this low-dimensional population regime is coordinated across hippocampal subfields. Correlations between dominant population trajectories (PC1 time series) revealed significantly stronger coupling between CA3–CA1, CA1–DG, and CA3–DG in enriched animals (**Fig. 5e**). Alignment of dominant population modes across regions has been proposed as a mechanism for large-scale integration of neural activity and effective inter-regional communication^20,59^, suggesting that enrichment promotes coordinated population dynamics across the hippocampal circuit. The enhanced cross-subfield alignment observed here provides a population-level substrate that links the expanded interaction geometry (Fig. 3) to the increased propagation of ripple events (**Fig. 4**).

To aid interpretation of these population-level analyses, we include a minimal schematic summary of the observed dynamical regime (**Fig. 5f**). This schematic distills the quantitative results by contrasting the diffuse, higher-dimensional population trajectories observed in standard-housed animals with the low-dimensional, PC1-dominated population dynamics in enriched animals. Importantly, the schematic does not introduce additional analyses but provides a conceptual synthesis linking the structured state-space trajectories (**Fig. 5a**), reduced dimensionality and increased PC1 dominance (**Fig. 5b–c**), enhanced population recruitment (**Fig. 5d**), and strengthened inter-regional coordination (**Fig. 5e**) into a unified population-dynamical framework.

Together, these results show that environmental enrichment reorganizes hippocampal population activity into a strongly coordinated, low-dimensional regime characterized by dominant shared modes, broad neuronal recruitment, and cross-subfield alignment. This population-level compression provides a unifying dynamical signature linking enrichment-dependent changes in basal activity, interaction geometry, and ripple organization.

## Discussion

This study demonstrates that sustained experience reorganizes hippocampal computation by showing that its effects are expressed across multiple organizational levels—basal network state, interaction geometry, event structure, and population dynamics—when measured simultaneously in the intact brain. Rather than producing isolated changes in signal amplitude or event rate, experience restructures how activity is distributed, aligned, and propagated across hippocampal subfields, thereby limiting the space of accessible network dynamics^4,60,61^.

A central implication of these findings is that experience-dependent plasticity cannot be fully understood at the level of local synaptic or cellular changes alone^2,12,13^. While such mechanisms are necessary substrates, they do not specify how computation unfolds across interacting populations. Our results instead support a circuit-level view in which experience reconfigures the geometry of interactions, changing the balance between local activity, long-range coordination, and population-level organization. This view aligns with theoretical and empirical work showing that correlation structure, interaction topology, and dimensionality place fundamental limits on information flow and representational capacity in neural systems^55,62,63^.

Notably, the reorganization observed here spans multiple dynamical scales that are typically studied in isolation. Experience-dependent changes in basal activity statistics, functional interaction geometry, and ripple engagement converge at the level of population dynamics, manifesting as a coordinated, low-dimensional regime dominated by shared modes. Rather than reflecting a loss of complexity, such population-level compression indicates structured coordination, consistent with models in which effective computation arises within constrained subspaces shaped by circuit history and connectivity^64^. From this perspective, population dynamics provide a unifying description of how diverse experience-dependent effects are integrated at the circuit level.

Although hippocampal dynamics are strongly state-dependent^65–67^, the convergence of experience-related changes across basal activity statistics, interaction geometry, ripple organization, and population dimensionality—together with their consistency with prior large-scale *ex vivo* observations^15,16,68^—cannot be readily explained by a purely state-driven account. Beyond transient-state modulation, the observed reorganization is consistent with experience-dependent shaping of circuit interactions structure. Prior work has shown that long-term experience can reshape population coupling, coordination geometry, and the balance between shared and independent activity modes without altering gross firing rates or oscillation frequencies. Such constraints act at the level of network organization, influencing how activity propagates and stabilizes across populations rather than dictating moment-to-moment state^69–71^.

A critical aspect of this study is that these multiscale effects were resolved within the same recordings, only through dense, simultaneous electroanatomical sampling. Addressing experience-dependent circuit computation requires simultaneous access to distributed populations, laminar structure, and inter-regional interactions within the same recordings. Critically, this depends not only on channel count but on spatial geometry: parallel, multi-shank sampling that spans extended circuit axes. Without such geometry, dense measurements along a single trajectory or sequential recordings cannot resolve how activity is coordinated across hippocampal subfields and layers at the circuit scale, making it difficult to distinguish local modulation from reorganization of circuit-wide dynamics^72^.

This study examines environmental enrichment as a sustained experiential modulation within the hippocampal formation, serving as a tractable model for experience-dependent circuit reorganization. Other forms of experience—including learning, stress, or pathology—may shape circuit dynamics through different organizational regimes, which may be expressed differently across brain systems. The analytical framework established here provides a general approach for testing how diverse experiential histories reshape circuit-level dynamics in intact networks, while the extent to which similar multiscale reorganization applies beyond the hippocampus remains an open question.

More broadly, these results have implications for interpreting and interfacing with neural systems. Increasing evidence indicates that effective interaction with neural systems—whether for decoding, stimulation, or restoration of function—requires understanding how experience structures population dynamics and shapes the space of accessible neural states^33,34^. From this perspective, experience-dependent restructuring of population dynamics defines the operating regime of a circuit and determines how external inputs or interventions can engage it. This understanding is therefore essential for computational models of learning and memory, as well as for the design of closed-loop neural interfaces and neuromodulation strategies that aim to interact with circuits shaped by experiential history rather than abstracted from it^60,73–75^.

## Methods

### Animal Experiments and *in vivo* neural recordings

All procedures were conducted in accordance with European and national regulations (Tierschutzgesetz) and approved by the Landesdirektion Sachsen (license number 25-5131/609/62). Experiments were performed on 12-week-old female C57BL/6J mice (Charles River Laboratories, Germany). Animals were received at five weeks of age and randomly assigned at six weeks to either standard housing (SD) or enriched environment (ENR) housing conditions. Mice were anesthetized with fentanyl (0.05 mg kg⁻¹), midazolam (5 mg kg⁻¹), and medetomidine (1 mg kg⁻¹), supplemented with carprofen (5 mg kg⁻¹) for perioperative analgesia, and positioned in a stereotaxic frame. A craniotomy was performed above the left dorsal hippocampus (AP −2.5 mm, ML −3.5 mm). Body temperature was maintained at 37 °C, and heart rate and respiration were continuously monitored using a Marta Pad (Vigilitech AG, Germany). A 1024-channel, 8-shank active CMOS SiNAPS probe (Corticale Srl, Italy) was inserted along the dorsoventral axis to a depth of 2.6–3.2 mm. Each shank spanned the hippocampal formation, enabling simultaneous recording across CA1, CA3, and DG. A platinum wire placed in the frontal skull and contacting cerebrospinal fluid served as a reference electrode.

### Data Acquisition, Signal Processing, and Neural Events Extraction

Neural signals were acquired using a Plexon OmniPlex system (Plexon Inc., USA) configured for SiNAPS high-density probes via the Digital Headstage Processor (DHP). Analog neural signals were first amplified locally at the electrode level by in-pixel amplifiers integrated on the SiNAPS probe. Signals were then digitized at the headstage and transmitted digitally via shielded headstage cables to the DHP, minimizing analog signal degradation and cable-borne noise. The DHP performed synchronized digital acquisition across all channels and streamed wideband data to the OmniPlex chassis through a high-speed digital link. Signals were sampled at an effective rate of 40 kHz per channel, providing temporally aligned wideband recordings across the full 1024-channel array. Digital referencing and filtering were applied within OmniPlex at the full wideband sampling rate before source separation. All analyses were performed on continuous channel-level signals without spike sorting. Wideband recordings were processed using CorticaleXplorer (CortiX) (Corticale Srl, Italy) for signal preprocessing and event detection. LFPs were obtained by band-pass filtering (2–100 Hz), while MUA was extracted by high-pass filtering (>300 Hz) and detecting threshold-crossing events on each channel. This approach captures aggregate population spiking while preserving the spatial structure of activity across the high-density array. SWR events were detected from the ripple band (120–240 Hz) using custom detection routines applied to the filtered LFP signal, followed by validation across channels and shanks to assess their spatial extent and laminar organization^16,76^. All downstream analyses were based on these continuous, channel-level representations of LFP, MUA, and SWR activity.

### Electroanatomical Registration and Region Assignment

Recording channels were assigned to anatomical regions using post hoc atlas-based registration. Probe geometry (shank spacing and electrode depth) was aligned to the Mouse Brain Atlas^77^ using histological reconstruction and insertion depth measurements. Channels were assigned to CA1, CA3, DG based on dorsoventral position relative to region boundaries. All subsequent analyses were performed on region-labeled channel sets.

## Data Analysis

All data processing and analyses were performed using custom-written Python scripts developed for this study. Analyses combined standard scientific computing libraries with bespoke implementations tailored to large-scale, multi-channel electrophysiological data. Where third-party packages or add-on libraries were used, they are cited accordingly. All analyses were applied consistently across experimental conditions and animals using identical parameter settings unless explicitly stated otherwise.

### Basal Activity Statistics and Lognormal Dynamics

We characterized the statistical structure of spontaneous activity at the channel and regional levels computed from LFP time series and MUA event trains. For each channel, distributions of LFP rate and MUA event rate were constructed over the full recording duration and pooled within anatomical regions (CA1, CA3, DG). Additionally, following analytical frameworks to capture the heavy-tailed organization of channel-resolved neural activity established in our prior studies^16,30,78^, firing distributions were analyzed on logarithmic scales and modeled using a lognormal distribution,

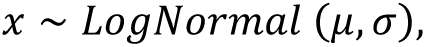

where 𝑥 denotes the channel-wise activity measure (LFP or MUA event rate), 𝜇 is the mean of the log-transformed channel activity, and 𝜎 is the mean of the standard deviation of the log-transformed channel activity. Outcome variables were region-resolved distributions of channel-wise activity and the fitted lognormal parameters 𝜇 and 𝜎.

### LFP–MUA Coupling Analysis

To quantify the relationship between mesoscopic network activity and population spiking output at the channel level, LFP and MUA firing rates were computed for each channel over the full recording duration. Coupling between LFP and MUA activity was quantified by assessing the dependence of MUA firing rate on LFP event rate across channels. Channel-wise coupling strength was estimated using linear regression applied to the joint distribution of LFP and MUA rates. Coupling metrics were summarized within anatomical regions and across the full hippocampal dataset. Outcome variables were channel-level LFP and MUA rates and region-averaged coupling coefficients.

### Functional Connectivity Analysis

To quantify large-scale statistical dependence between hippocampal recording sites across anatomical regions, pairwise functional connectivity was computed between all recorded channels using Pearson’s correlation coefficient. For each recording session, LFP signals and MUA activity were extracted for all 1,024 channels. For each channel pair (𝑖, 𝑗), functional connectivity was defined as 𝜌_ij_ = 𝑐𝑜𝑟𝑟(𝑥_i_, 𝑥_ij_), where 𝑥_i_and 𝑥_ij_ denote the LFP signal or MUA activity recorded from channels (𝑖, 𝑗), respectively. Correlations were computed over the full analysis window, yielding a symmetric 1,024×1,024 correlation matrix for each session and signal type. Channels were assigned to CA1, CA3, and DG based on atlas-aligned electrode localization, and Pearson correlation matrices were analyzed at the channel level and after averaging across within-and across-region channel pairs for LFP and MUA signals.

### Graph-Theoretic Network Analysis

To quantify topological organization and directional information flow in hippocampal functional connectivity, directed functional networks were constructed from pairwise Pearson correlation matrices computed from LFP and MUA signals. We computed the graph metrics in custom-written Python code and adapted functions from NetworkX-python package^79^, available on GitHub (https://github.com/networkx). Briefly, nodes corresponded to recording channels, and edges represented statistically dependent functional connections between channel pairs. Directional connectivity matrices were obtained by grouping correlation coefficients according to ordered anatomical region pairs (e.g., CA1→CA3, CA3→CA1), as previously implemented^15,16,78^. Adjacency matrices 𝐴 = {𝑎_ij_} were defined by thresholding correlation values, where 𝑎_ij_ denotes a directed functional connection from nodes 𝑖 to node 𝑗.

To characterize the different representations of network connectivity, we characterized the degree 𝑘 of a node 𝑛 to describe the number of edges connected to a node. Node degree was defined as

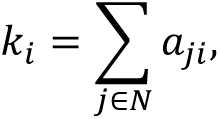

where 𝑁 denotes the set of all nodes.

To measure how nodes in a given network tend to cluster together to assess the functional segregation, the local clustering coefficient (𝐶𝐶_i_) was computed as

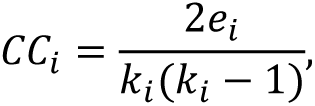

Where 𝑒_i_denotes the number of edges between theneighbors of node 𝑖. Node-level metrics were averaged across channels assigned to CA1, CA3, and DG.

### Spatial Interaction Scale Analysis

This analysis quantified the spatial extent of functional correlation between hippocampal LFP and population spiking activity. For each recording session, pairwise Pearson correlation coefficients were computed between all electrode pairs using LFP signals and, separately, MUA activity. For each electrode pair (𝑖, 𝑗), the physical distance 𝑑_ij_ was calculated from the known probe geometry using shank index and depth coordinates. Correlation coefficients were grouped into non-overlapping distance bins (e.g., 0–100 µm, 100–200 µm). For each distance bin 𝑏, mean functional connectivity was computed by averaging correlation values across all electrode pairs whose inter-electrode distance fell within that bin. This procedure was applied separately for LFP-and MUA-based connectivity. Electrode pairs were further classified as within-region or across-region based on atlas-aligned anatomical assignments (CA1, CA3, DG), and distance-dependent correlation profiles were computed independently for each class. Correlation–distance curves were generated separately for SD and ENR conditions. Outcome variables consisted of mean Pearson correlation values as a function of inter-electrode distance for LFP and MUA signals, stratified by within-and across-region electrode pairs and experimental condition.

### LFP–Spike Correspondence of Functional Connectivity

This analysis quantified the correspondence between mesoscopic field-based and population spiking–based functional connectivity across hippocampal regions. For each recording session, region-pair functional connectivity was computed separately for LFP and MUA signals using Pearson correlation coefficients. For a given region pair (e.g., CA3–CA1, CA1–DG, CA3–DG), LFP connectivity was defined as the mean Pearson correlation across all channel pairs spanning the two regions, and spike connectivity was defined analogously using MUA correlations. For each animal and region pair, LFP-based connectivity values were paired with the corresponding spike-based connectivity values. Analyses were performed separately for SD and ENR animals. Outcome variables consisted of paired LFP and MUA region-level connectivity values for each animal and region pair, used for cross-modal comparison of functional interactions.

### Ripple Detection and Event Characterization

SWR events were detected to identify transient, high-frequency hippocampal network events and quantify their temporal and amplitude properties, using a previously described approach^16,39^. Briefly, whiteband signals were band-pass filtered in the ripple band (120–240 Hz), and the analytic amplitude was obtained using a Hilbert transform and smoothed with a Gaussian kernel (σ = 4 ms). Channel-level ripple segments were identified when the smoothed envelope exceeded 3 standard deviations above baseline for a minimum duration of 15 ms. Channel-level detections were merged into global ripple events based on temporal overlap, as previously implemented. Ripple rate was defined as the number of detected global events per minute. Ripple amplitude was defined as the peak ripple-band envelope within each event, and event duration as the interval between threshold crossings. Outcome variables included ripple rate, peak amplitude, and inter-event duration, computed per region and recording session, and compared across conditions.

### Ripple Spatial Recruitment and Propagation

This analysis quantified whether detected ripple events remained local to a single subfield or co-engaged CA3 and CA1. For each detected ripple event, all channels exhibiting ripple-band activity during the event window were identified and assigned to hippocampal regions (CA1, CA3) based on atlas-aligned electrode localization. Channel-level detections were consolidated into global ripple events, and each event 𝑒 was assigned the set of anatomically engaged regions.

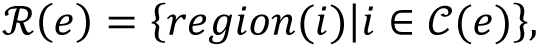

Where 𝒞(𝑒) is the set of channels exhibiting ripple-band threshold crossing during event 𝑒, and 𝑟𝑒𝑔𝑖𝑜𝑛(𝑖) ∈ {𝐶𝐴1, 𝐶𝐴3}, is the atlas-aligned assignment of channel 𝑖. Events were classified as:

- CA1-only if ℛ(𝑒) = {𝐶𝐴1}
- CA3-only if ℛ(𝑒) = {𝐶𝐴3}
- CA3-CA1 co-engaged if {𝐶𝐴1, 𝐶𝐴3} ⊆ ℛ(𝑒)

The Ripple Recruitment Index (RRI) was defined as the fraction of CA1/CA3 ripples that co-engaged both subfields relative to those remaining local:

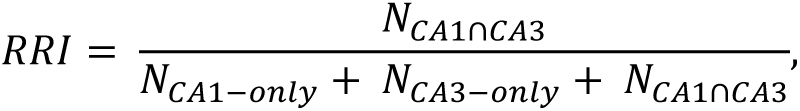

where 𝑁_CA1∩CA3_ is the number of events with CA1–CA3 co-engagement, and 𝑁_CA1-only_, 𝑁_CA3-only_ are the numbers of events confined to CA1 or CA3, respectively. Outcome variables included the proportions of local and co-recruited ripple events and the RRI, computed per animal and compared across experimental conditions.

### Ripple-triggered Broadband LFP

This analysis quantified the laminar LFP response time-locked to ripple events across depth and region. For each global ripple event 𝑘 a reference time 𝑡_O,k_was defined from the ripple detector (ripple-band envelope peak in a canonical CA1 pyramidal-layer channel, or an equivalent fixed reference rule applied to all sessions). Broadband 𝐿𝐹𝑃_ch_(𝑡) (2–100 Hz; not ripple-band filtered) was segmented into event-centered snippets for every channel 𝑐ℎ:

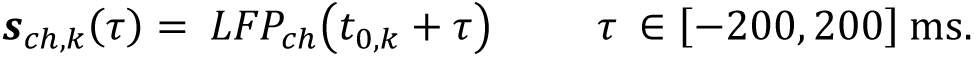

Snippets were averaged across detected ripples within each condition to obtain a ripple-triggered LFP for each channel:

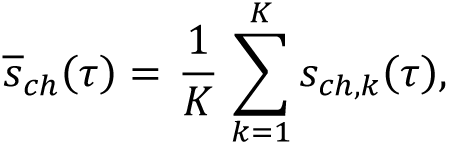

where 𝑘 is the number of ripple events used for that condition. Channels were ordered by probe geometry (shank × depth) and anatomical assignment (CA1/CA3) for laminar maps of 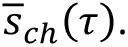 Peak ripple-triggered LFP amplitude was computed per region from the channel-wise 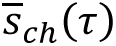 as the extremum within a fixed post-trigger window (same window across conditions), and peak latency was defined as the time of that extremum relative to 𝜏 = 0 (ripple reference time). Outcome variables were 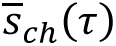 (channel-wise ripple-triggered broadband LFP), region-level peak amplitude, and region-level peak latency, computed per animal and condition.

### Population State-Space Analysis and Dimensionality Metrics

The analysis was performed to characterize the low-dimensional structure and dominant modes of hippocampal population spiking activity. For each recording session, population activity matrices were constructed by binning multiunit spiking activity across all recorded channels into consecutive time bins, yielding a matrix 𝛸 ∈ ℝ^fxN^, where 𝑇 is the number of time bins, and 𝑁 is the number of channels. Spike counts were z-scored per channel across time to remove rate offsets.

Principal component analysis (PCA) was applied to 𝛸 to identify dominant population activity patterns. Time-resolved population trajectories were obtained by projecting each time bin onto the leading principal components, generating a low-dimensional representation of population dynamics.

Population dimensionality was quantified using the participation ratio (PR) computed from the PCA eigenvalue spectrum {𝜆_i_},

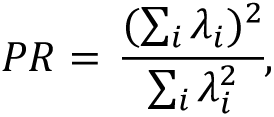

which provides an effective dimensionality measure of population activity. In addition, variance fractions explained by individual principal components were computed, and PC1 dominance was defined as the fraction of total variance explained by the first principal component. Outcome variables consisted of state-space trajectories, participation ratio, principal component variance fractions, and PC1 dominance, computed per animal and condition.

### Population Recruitment Metrics

Population recruitment analysis was performed to quantify the extent and variability of channel-level participation in ongoing hippocampal activity. For each time bin, the number of active channels was computed by counting channels exhibiting at least one spike within the bin. This yielded a time series of population recruitment values describing the instantaneous size of the active population. Recruitment metrics were computed across all channels and separately within anatomical regions (CA1, CA3, DG) based on atlas-aligned channel assignments. Outcome variables consisted of the distribution and summary statistics of active-channel counts per time bin, computed globally and region-resolved for each animal and condition.

### Inter-regional Population Coordination

To quantify alignment in population dynamics across hippocampal subfields, region-specific population activity matrices were constructed by restricting the full population matrix to channels within each anatomical region. PCA was performed separately for each region, and time-resolved trajectories were obtained in the corresponding low-dimensional spaces. Inter-regional coordination was quantified by computing correlations between region-specific population trajectories in latent space, using matched principal component dimensions and aligned time bins. Outcome variables consisted of cross-region latent-space correlation values, computed per animal and condition.

## Statistical Analysis

All were performed to evaluate differences between SD and ENR groups across all reported measures. Analyses were conducted on n=4 animals per group. All data in this work were expressed as the mean ± standard error of the mean (SEM). All box charts are determined by the 25th-75th percentiles and the whiskers by the 5th-95th percentiles and lengths within the Interquartile range (1.5 IQR). Also, the lines depict the median and the triangle for the mean values. Group comparisons of summary statistics were performed using one-way analysis of variance (*ANOVA*), or *two-way ANOVA* followed by Tukey’s *post hoc* testing, whereas comparisons of full distributions were assessed using the two-sample *Kolmogorov–Smirnov test*. Multiple comparisons arising from region-resolved or metric-wise analyses were controlled within each analysis family. All statistical tests were two-sided, and statistical significance was defined as P<0.05.

## Authors Contributions

S.K. performed experiments, analyzed data, and generated figures. X.H. contributed to data analysis and code implementation. D.K. assisted with animal experiments. G.K. conducted environmental enrichment procedures. F.B. contributed to data interpretation and discussion. H.A. conceived and supervised the study, designed and performed experiments, developed computational and analytical frameworks, conducted multiscale data analysis, and generated figures. All authors reviewed and approved the final manuscript.

## Acknowledgments

This work was supported by institutional funding from the Deutsches Zentrum für Neurodegenerative Erkrankungen (DZNE). We thank the behavioral animal testing platform at DZNE Dresden for technical support, including Alexander Garthe, Anne Karasinsky, Jens Bergmann, and Sandra Günther. We acknowledge Matteo Falappa (Corticale Srl, Italy) for helping with data processing using CortiX. We also thank Harvey Wiggins, Craig Patten, and Damon Gee (Plexon Inc., USA) for valuable technical discussions.

